# Influence network model uncovers relations between biological processes and mutational signatures

**DOI:** 10.1101/2021.11.16.468828

**Authors:** Bayarbaatar Amgalan, Damian Wojtowicz, Yoo-Ah Kim, Teresa M. Przytycka

## Abstract

There has been a growing appreciation recently that mutagenic processes can be studied through the lenses of mutational signatures, which represent characteristic mutation patterns attributed to individual mutagens. However, the causal link between mutagens and observed mutation patterns remains not fully understood, limiting the utility of mutational signatures. To gain insights into these relationships, we developed a network-based method, named GeneSigNet that constructs a directed network among genes and mutational signatures. The approach leverages a sparse partial correlation among other statistical techniques to uncover dominant influence relations between the activities of network nodes. Applying GeneSigNet to cancer data sets, we uncovered important relations between mutational signatures and several cellular processes that can shed light on cancer related mutagenic processes. Our results are consistent with previous findings such as the impact of homologous recombination deficiency on a clustered APOBEC mutations in breast cancer. The network identified by GeneSigNet also suggest an interaction between APOBEC hypermutation and activation of regulatory T Cells (Tregs) and a relation between APOBEC mutations and changes in DNA conformation. GeneSigNet also exposed a possible link between the SBS8 signature of unknown aetiology and the nucleotide excision repair pathway. GeneSigNet provides a new and powerful method to reveal the relation between mutational signatures and gene expression.

GeneSigNet is freely available at https://github.com/ncbi/GeneSigNet.

## 1 Introduction

Traditionally, research in cancer genomics has been focused on the identification of cancer driving mutations that confer a growth advantage to cancer cells. However, since cancer often emerge as a byproduct of various mutagenic processes such as UV light or a faulty DNA repair mechanism, cancer genomes also accumulate numerous mutations with seemingly no direct roles in carcinogenesis. Different mutagenic processes leave different patterns of somatic mutations called *mutational signatures*. In addition, different cancer genomes can be subjected to different levels of exposures to such mutagenic processes. The exposure of a genome to a given mutagenic process is measured by the number of mutations attributed to this process.

Starting from the pioneering work of Alexandrov et al. [1], several computational methods have been developed to infer mutational signatures. However, these methods focus on defining mutational signatures as patterns of mutations but do not link these patterns to specific mutational processes. The relations between these computationally derived signatures and mutational processes causing them were established based on prior knowledge, association between mutational processes and environmental factors or dedicated experimental studies. In this way, several mutational signatures have been linked to specific mutagenic processes [2,3]. However, the etiology of many signatures still remains unknown or not fully understood. At the same time computational approaches that can assist in the identification of relations between mutational signatures and cellular processes have been extremely limited.

Elucidating mutational signatures associated with mutagenic processes is of fundamental importance for a better understanding of carcinogenesis and for facilitating the development of novel treatments. Gene expression provides the most accessible measurement of the activities of cellular processes. Therefore, we reasoned that interrogating the relation between gene expression and exposure of a mutational signature might provide important clues on the etiology of the signature. For example, deficiencies in the activities of several genes such as MUTYH [4], ERCC2 [5], MSH6 [6], and FHIT [7] have been linked to specific signatures. In addition, a correlation between the expression of the APOBEC family of genes and the exposures of signatures SBS2 and SBS13 (so-called APOBEC mutational signatures) has frequently been observed [2,8]. However, the relations between mutational signatures and gene expression can be quite complex. For instance, tobacco smoking is not only mutagenic itself but it is also believed to activate the immune response [9]. In addition, a perturbation of some cellular processes such as DNA replication or repair pathways can also be mutagenic. Furthermore, mutagenic processes themselves have been known to interact with each other: homologous recombination deficiency (HRD) is typically accompanied by a mutational signature related to APOBEC activity [10,11]. Applying a clustering-based method to genome-wide expression and mutational signature data, Kim et al. identified coherently expressed groups of genes associated with specific combinations of mutational signatures providing interesting clues about these signatures [11]. However, such cluster-based analysis only suggest broadly defined associations between signature exposures and the activities of biological processes, lacking more precise and mechanistic explanations.

To fill this gap, we introduce a network-based method, named GeneSigNet (Gene and Signature Influence Network Model), aiming to uncover interactions among mutagenic and cellular processes, and to provide insights into mutagenic processes underlying individual signatures. Utilizing gene expression and mutational signature data from cancer patients, GeneSigNet constructs a GeneSignature Network (GSN) that provides directed relationships among two types of node entities – genes and MutStates. MutState nodes are in one-to-one correspondence with mutational signatures and each MutState represents an abstract (directly unobserved) cell state associated with the emergence of the corresponding mutational signature (see Section 2.1 for a detailed description). Both types of nodes have patient-specific activities: gene expression for nodes corresponding to genes and signature exposure for nodes corresponding to MutStates. GeneSigNet utilizes these activities to infer direct edges between nodes in GSN (Fig. 1).

**Fig. 1.**
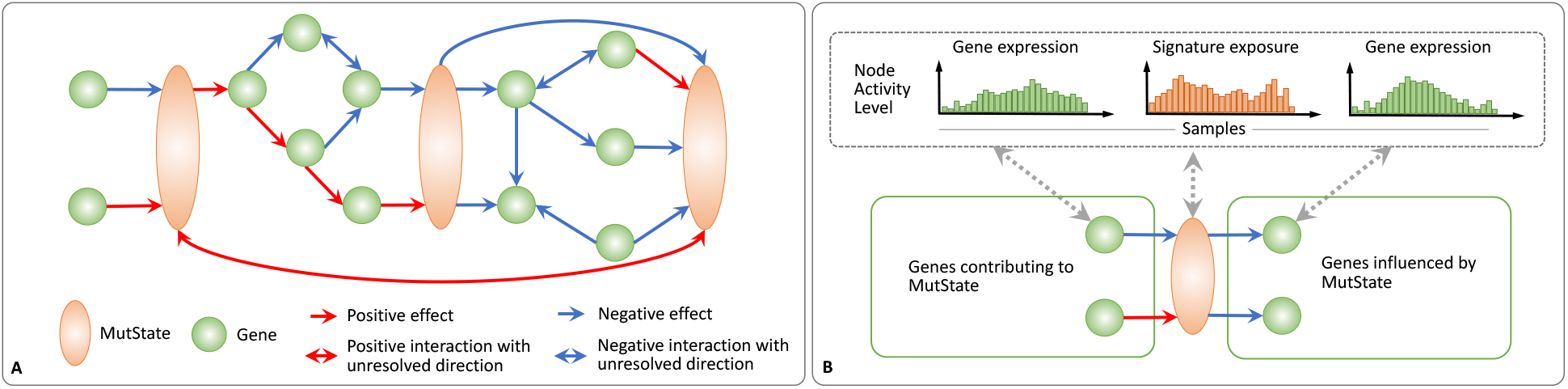
Gene-Signature Network (GSN). **(A)** GSN is a weighted-directed network consisting of two types of nodes: genes (green circles), and MutStates (orange ovals) corresponding to signatures. Edges of a GSN represent inferred influences and might be either positive (red) or negative (blue). **(B)** Edges of a GSN are inferred based on the activities of nodes (gene expression for genes and signature exposures for MutStates (top panel)). A directed edge from a gene to a MutState represents a putative influence from the gene to the MutState. Analogously, a directed edge from a MutState to a gene represents an inferred effect of the MutState on the gene.

Ideally, we would like to infer causal relationships between molecular activities and mutational signatures. However, in complex molecular systems, inferring causality is one of the most challenging open problems. In some settings, a prior information about the gene function (such as transcription factor versus potential target gene) or special type of data such as response to a perturbation of specific genes can be leveraged to infer direction [12,13,14,15]. In our setting, however, no perturbation data or prior knowledge is available, requiring GeneSigNet to rely solely on the observed activities of nodes for inferring edges and their directions. While inferring directed influences without such additional information remains challenging, significant efforts have been made to identify the conditions where such information can be inferred with a reasonable level of success. For example, independent component analysis-based method LiNGAM allows to discover the causal relations between non-Gaussian random variables when the number of variables is much smaller than the number of observations [16]. Partial correlation-based heuristic GeneNet constructs a directed acyclic causal network as a subgraph of partial correlation network based on a multiple testing of standardized partial variances [17]. A sparse estimation of partial correlations (SPCS) computes sparse asymmetric weight matrix representing directed influences among nodes in the network. Due to its sparsity constraints many entries of such inferred matrix are equal to zero and thus some pairs of nodes will be connected by one-directional edge only while a significant fraction of the edges have nonzero weights in both direction [18]. In contrast, the statistical higher moments were used as indicators of the direction of dependency between two variables [19] under the assumption of absence of confounding effects. Building on the last two approaches, GeneSigNet focuses on inferring most informative relations by performing two complementary steps. First, a sparse partial correlation technique (SPCS) is used to obtain an initial sparse weighted directed network. This graph contains both unidirectional and bidirectional edges. Next, where applicable, a partial higher moment strategy is used to resolve bidirectional edges. We note that GeneSigNet does not attempt to construct a fully resolved directed graph leaving many edges as bidirectional. We found that this approach compares favorably to the previously proposed techniques for this task.

We applied GeneSigNet to two cancer datasets, breast cancer and lung adenocarcinoma, for which sufficient numbers of patient samples with gene expression data are available and the interactions of mutational signatures are partially known. The relations inferred by the GeneSigNet model are overall consistent with current knowledge but also include several interesting novel findings. In particular, the model suggests a causative relation from the homologous recombination deficiency signature (SBS3) to a clustered APOBEC mutation signature, and also linked Signature 8 (SBS8) to nucleotide excision repair (NER) pathway. The latter connection is consistent with the recent findings based on an experimental study in mouse [20]. In addition, GeneSigNet identified a novel relation between APOBEC hypermutation and the activation of regulatory T cells which presents an important implication in immunotherapy, and captured a relation of APOBEC signature (SBS2) with DNA conformation changes among other findings. Our results demonstrate that GeneSigNet provides novel and important insights that were not accessible with existing methods.

## 2 Results

### 2.1 The GeneSigNet method

#### Gene-Signature Network

The main idea behind GeneSigNet is a construction of a Gene-Signature Network (GSN) consisting of two types of nodes: nodes corresponding to genes and nodes corresponding to mutational signatures (Fig. 1A). Patient-specific activity of a node corresponding to a gene is measured by gene expression. The exposure of a mutational signature can be seen as a measure of the activity of the corresponding mutagenic process, such as an exogenous mutagen, or the activity of a cellular process triggered in the response to a mutagen. This motivates the concept of *MutState* defined as an abstract representation of cellular state associated with a specific mutational signature. The activity of this state is measured as the exposure of the corresponding mutational signature. Statistical relations on the activities of genes and MutStates are likely to shed light on the genes and pathways associated with and potentially contributing to the level of the exposure of the corresponding mutational signature. To this end, GeneSigNet infers directed edges between both types of nodes, utilizing patient-specific gene expression (for genes) and exposures of mutational signatures (for MutStates) as described below (Fig. 1B). Other than the difference in the definitions of node activities (and subsequent interpretation), the network inference algorithm does not distinguish between the two types of nodes.

#### A high level description of the GeneSigNet inference method

GeneSigNet method constructs GSN in two main steps: (i) constructing a preliminary directed network using a sparse partial correlation selection (SPCS) and (ii) revising the initial network by testing whether the directionality of bidirectional edges can be decided using a partial higher moment strategy adapted to a network setting. The workflow of the GeneSigNet method is shown in Fig. 2.

**Fig. 2.**
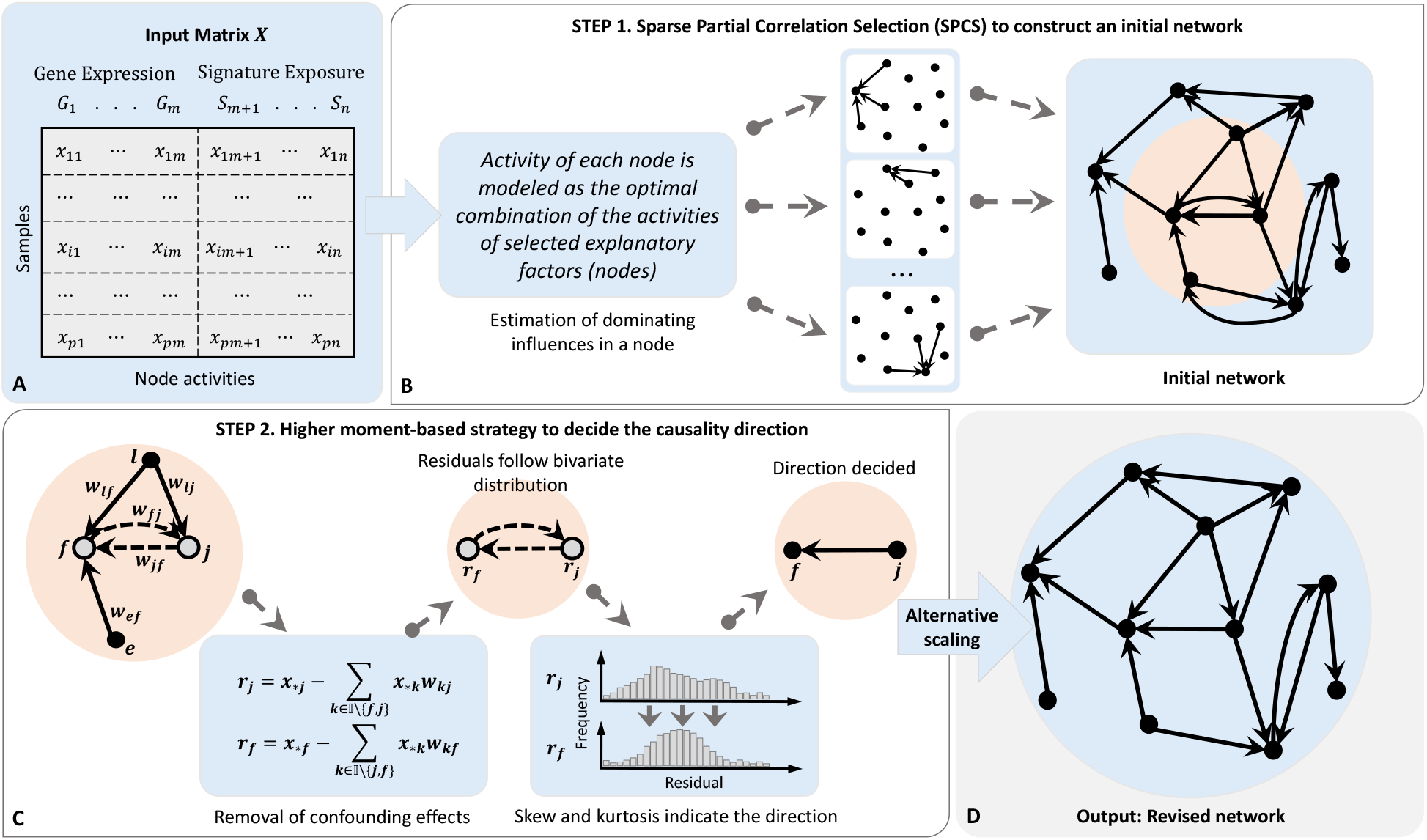
Workflow of GeneSigNet. **(A)** Input matrix *X* of node activities constructed by concatenating gene expression values (for genes) and signature exposures (for MutStates) across *p* samples (patients). **(B)** Given the input matrix *X*, we infer a network of *n* nodes. For each node, Sparse Partial Correlation Selection (SPCS) is used to simultaneously estimate the weights of incoming effects from the other *n*− 1 nodes. **(C)** For each bidirectional edge (*j, f*), the residual vectors *r*_*j*_ and *r*_*f*_ corresponding to nodes *j* and *f* are obtained by removing effects of the *n*− 2 nodes other than the two nodes of the considered edge. The non-zero weight *w*_*kf*_ obtained by SPCS denotes the strength of the confounding effect on node *f* coming from node *k*. The direction of a influence effect between the pair of nodes is determined based on the partial higher moment statistics, skewness and kurtosis of residuals *r*_*j*_ and *r*_*f*_. If both moments support the same direction with the heavier partial correlation weight (see Equation S5 in Supplementary Information), then the edge corresponding to the opposite direction is removed, otherwise, both edges remain in the network. **(D)** Edge weights are normalized using an alternative scaling algorithm, and the final weighted-directed network is obtained as the output.

In the first step, given the input matrix describing the activities of genes and MutStates across samples, GeneSigNet constructs an initial weighted-directed graph using a sparse partial correlation selection (SPCS) (Fig. 2B). Specifically, considering each node as a target, GeneSigNet uses SPCS to compute the weights of incoming effects from the other *n* − 1 nodes. This SPCS step builds on the fact that partial correlations can be approximated by sparse regression coefficients (see Supplementary Sections S1.1 and S1.2 for a detailed description). We note that since SPCS is applied to each node separately, GeneSigNet ensures a locally sparse solution rather than constraining the network edges globally.

In the second step, GeneSigNet refines the initial network by reducing the number of bidirectional edges remaining after the first step. The idea is an adaptation of the basic bivariate higher moment strategy to multivariate analysis. Specifically, for any two nodes having potential effects on each other in the initial network (endpoints of a bidirectional edge), we first utilize the partial correlation technique to remove confounding effects due to the presence of the other *n* −2 variables from the observed activities of the two nodes and obtain the residuals representing the remaining dependencies between the pair [21]. Under the assumption that all confounding effects due to the presence of the other *n* −2 variables were successfully removed by partial correlation, the influence variable may be distinguished from the affected variable by comparing the higher moments of the two residual distributions. Specifically, the affected variable is expected to be closer to normality than the influence factor, and the skewness and kurtosis are the higher moment statistics used to measure the close-normality of distributions. We refer to this strategy as *partial higher moment strategy*.

We note that the method relies on simplifying assumptions which might not be fully satisfied in real biologic relations. In particular, the assumption that all confounding effects come from the activities of remaining *n* −2 nodes in the network is an oversimplification since in such complex systems, potential effects from unobserved latent factors are likely to be present. Thus, we next evaluated the performance of the method on simulated and real data (to the extend possible in the absence of the ground truth).

Fig. 3 provides a real example illustrating the necessity of removing confounding effects before applying the higher moment statistics to correctly indicate the influence direction. Before removing confounding effects, the two higher moment statistics provided contradictory

**Fig. 3.**
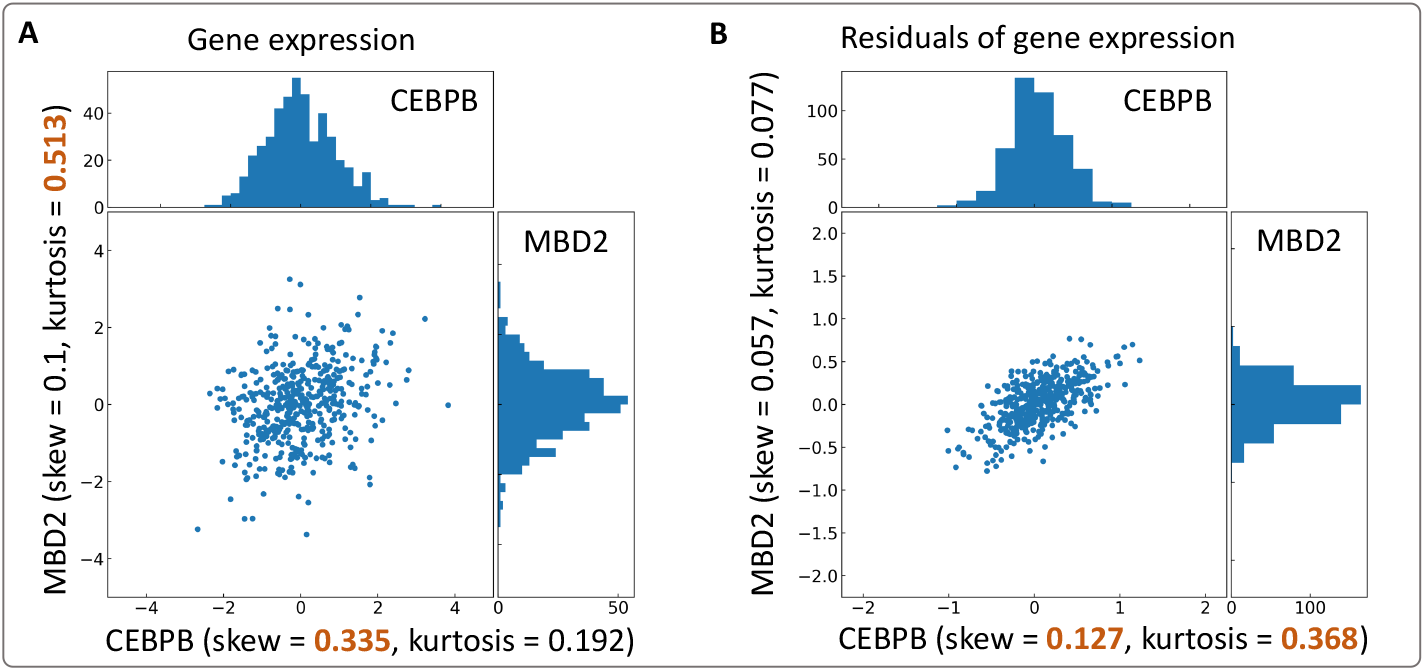
Basic higher moment statistics (A) versus the partial higher moment strategy (B). The experimentally confirmed regulatory relation is from CEBPB to MBD2. Using uncorrected moment statistics, the two higher moment statistics, skewness and kurtosis, provide contradictory directions (A). The proposed partial higher-moment strategy predicts the correct direction (B). For each case, brown color indicates the higher value for a given moment.

After removing the confounding effects, both statistics correctly indicate the influence direction [22] (Fig. 3B). More examples based on experimentally confirmed regulatory directions [22] are provided in Supplementary Figures Fig. S2.

Finally, a matrix normalization algorithm, alternate scaling [23], is used to bring the total incoming and outgoing effects of each node to the same range (see Supplementary Section S1.4 for details). A detailed description of GeneSigNet is provided in Supplementary Section S1.

#### Evaluation of the directionality inference on simulated data

We compared the performance of GeneSigNet to three competing approaches: (1) the independent component analysis-based method LiNGAM [16], (2) the partial correlation-based heuristic GeneNet [17] previously proposed to discover the causal structure in high-dimensional genomic data [18], and (3) the Sparse Partial Correlation Selection (SPCS) approach [18]. We used the SPCS method as the preliminary procedure to the partial higher moment strategy, which is described in Section 2.1. For completeness, we also included the regression tree-based approach used by GENIE3 method [24] for the inference of Gene Regulatory Networks (GRN) [24,25,26]. The task of GRN inference is quite different from inferring influence graph considered in this study. However, since the method produces a fully connected directed weighed graph we used a procedure that reduces bidirectional edges to directed edges by retaining the heavier edge in the pair.

To generate data, we implemented the data simulation schema provided in LiNGAM [16]. This schema starts with constructing a lower triangular weight matrix representing the directed interactions among the nodes (random variables) in a directed acyclic graph (DAG). This matrix is then used as the representation of the dependency patterns in the generation of the data set (see Supplementary Section S2.3 for detailed description of data generation). We simulated a set for 100 random variables with 1000 samples from multivariate distributions. The weight matrix used in the simulation contains 364 directed edges.

The performances are summarised in the Fig. 4 and the corresponding F-scores are provided in Fig. S3A in Supplementary Information. In addition to the evaluation for the true positive fraction of predicted edges, the correlation between the generated and estimated weight matrices is computed to show the reproducibility of influence weight values in the reference lower triangular matrix (Fig. S3B in Supplementary Information). GeneSigNet performed competitive with GeneNet on simulated data. The precision for the GeneSigNet prediction 0.707 (TN: 0.9903) whereas those are 0.59 (TN: 0.9412) for GeneNet and 0.560 (TN: 0.9923) for LiNGAM.

**Fig. 4.**
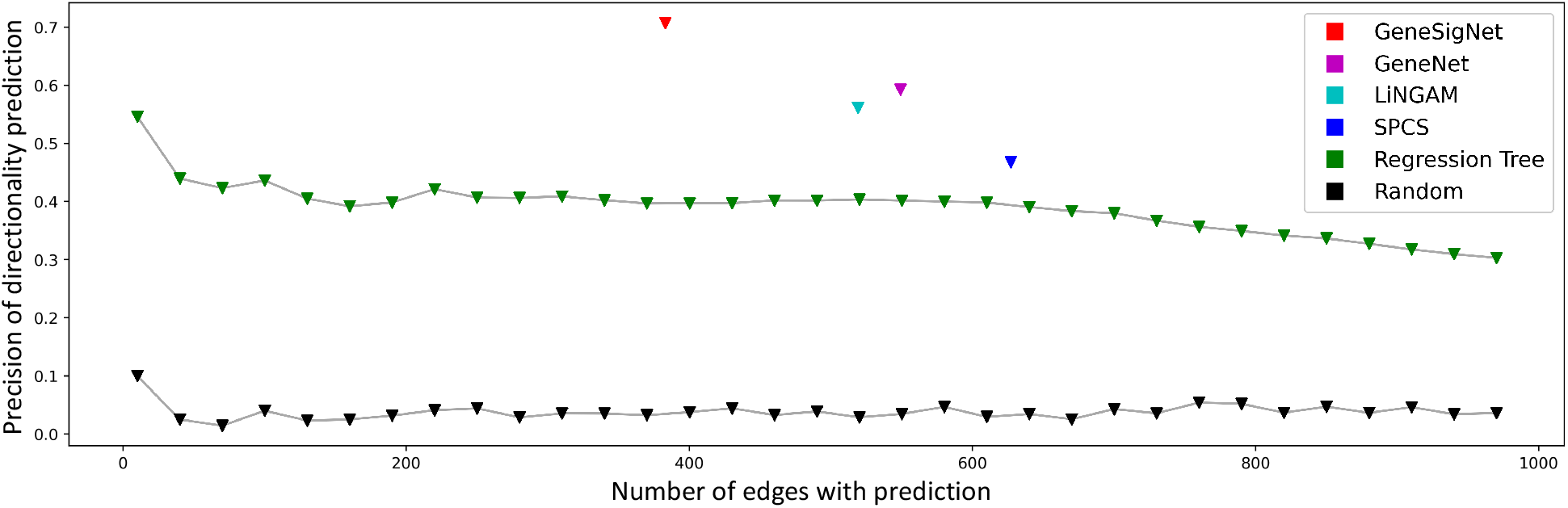
The horizontal axis denotes the number of selected edges in the prediction. The vertical axis denotes the precision of the directionality inference.

#### Evaluation of the directionality inference on real cancer data

As discussed above, in biological systems the assumption that all confounding effect is due to the activities of the remaining *n* −2 nodes and could be removed using partial correlation is an oversimplification and it is important to evaluate the method on real data. We compared the performance of the methods on breast cancer (BRCA) and lung cancer (LUAD) data sets (see Supplementary Section S2 for details). The LiNGAM method was excluded from the evaluation, since it cannot deal with the large genome-wide datasets.

To evaluate the accuracy of the directionality inference, we used the ChEA database containing directed protein-DNA interactions [22]. Specifically, as the gold standard set we used the set of one-directional edges that are in the intersection of ChEA and the network inferred by a tested method (Fig. 5).

**Fig. 5.**
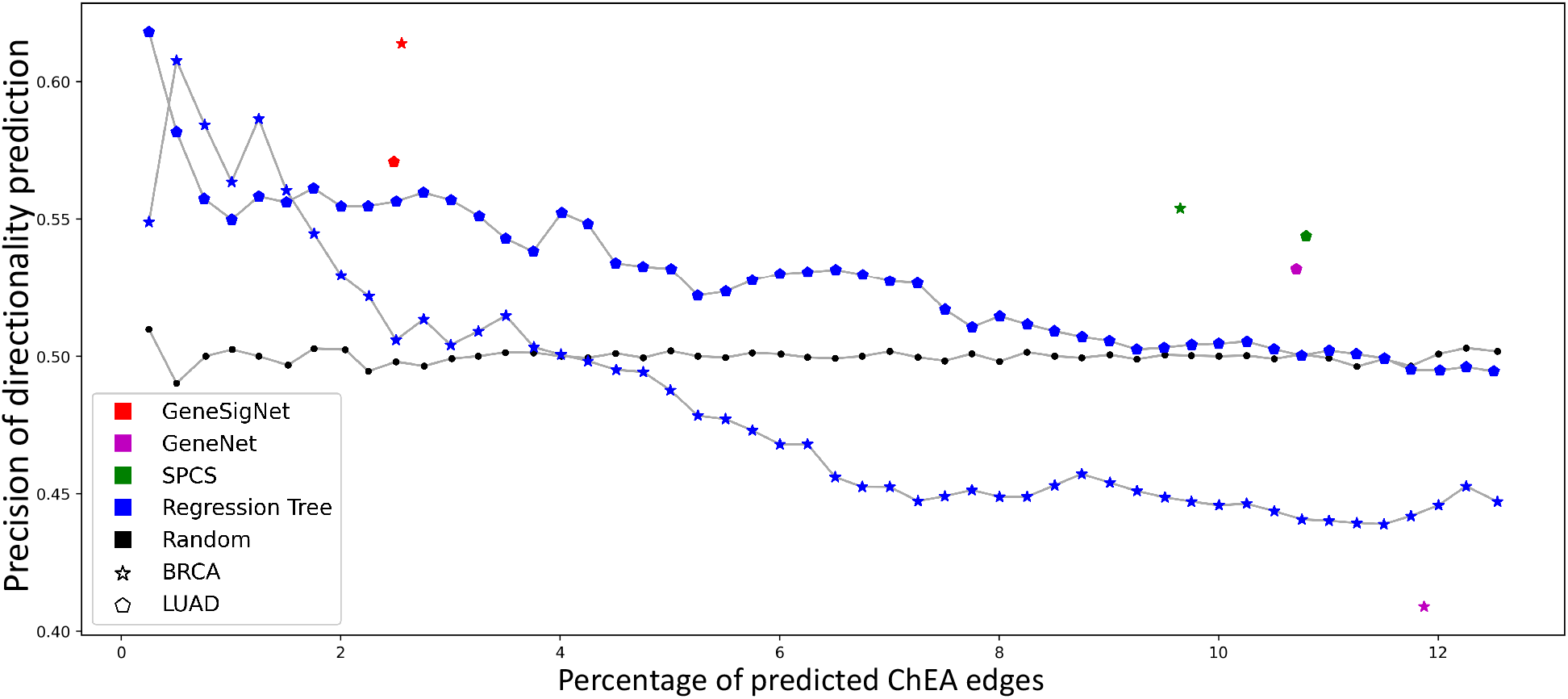
The evaluation of directionality inference on real data sets. The horizontal axis denotes the percent of predicted one-directional edges from ChEA data whereas the vertical axis denotes the fraction of correctly predicted ChEA directions.

In addition, we tested the robustness of the partial higher moment strategy by performing a bootstrap sampling which is described in Supplementary Section S1.3. The approach was highly reproducible as summarized in Section S1.3 and Fig. S1 in Supplementary Information. GeneSigNet compared favorably to the competing approaches. However, the performance of all the methods on the task of inferring correct direction of information flow was weaker on the real data than on simulated data.

### 2.2 Analysis of the relations between mutational signatures and molecular pathways in breast cancer

We utilized breast cancer (BRCA) data collection obtained from ICGC which includes 266 cancer samples providing both whole-genome sequencing data and gene expression data (for details, see Supplementary Section S2.1). The breast cancer genomes harbor mutations mainly contributed by 6 COSMIC mutational signatures – SBS1, 2, 3, 5, 8, and 13. We further refined the mutational signatures based on mutation density and sample correlations. The mutations in BRCA are characterized by occurrences of short highly mutated regions whose origin is believed to be different than sparse mutations [8,11,27,28,29]. The information available from whole-genome sequencing allows for distinguishing these two types of mutation patterns and to treat such dense and sparse mutation regions differently. The post-processing of mutational signatures resulted in 6 signature groups that we use for subsequent analysis to construct the GSN – SBS1, APOBEC-C (clustered SBS2 and SBS13 corresponding to APOBEC hypermutation), APOBEC-D (SBS2 corresponding to disperse APOBEC mutations), DSB (SBS3 and clustered SBS8), SBS5, and SBS8D (dispersed SBS8). In addition to gene expressions and exposures of mutational signatures, we included a node indicating the binary status of homologous recombination deficiency (HRD) as it is assumed to lead to specific patterns of mutational signatures in BRCA [30]. We applied GeneSigNet to construct a GSN for genes, mutational signatures, and HRD status, and to find relations between these features.

#### Consistency of GeneSigNet results with current knowledge

Many relations uncovered with GeneSigNet are consistent with our current knowledge on mutational signatures, confirming the validity of our method. In particular, it is well appreciated that homologous recombination (HR) plays an important role in the double-strand break (DSB) repair mechanism and that HR deficiency is associated with the DSB signature [31]. Indeed, our network correctly predicted a strong positive influence from HRD status to the DSB signature (Fig. 6). In addition, GeneSigNet identified the known negative impact of BRCA1 expression on the DSB signature which is also consistent with the role of BRCA1 in HRD [31]. Furthermore, GeneSigNet captured the impact of HRD on chromosome separation, reflecting the role of homologous recombination in maintaining genomic stability [32,33], and identified the association of APOBEC-D with telomere maintenance, consistent with the well recognized role of APOBEC mutagenesis in replication [34,35].

Interestingly, our method linked SBS8 to the nucleotide excision repair (NER) pathway (Fig. 6). The etiology of this signature has remained unknown until a recent experimental study linked it to the NER pathway as well [20]. This demonstrates the power of the GeneSigNet method to uncover non-obvious relationships.

**Fig. 6.**
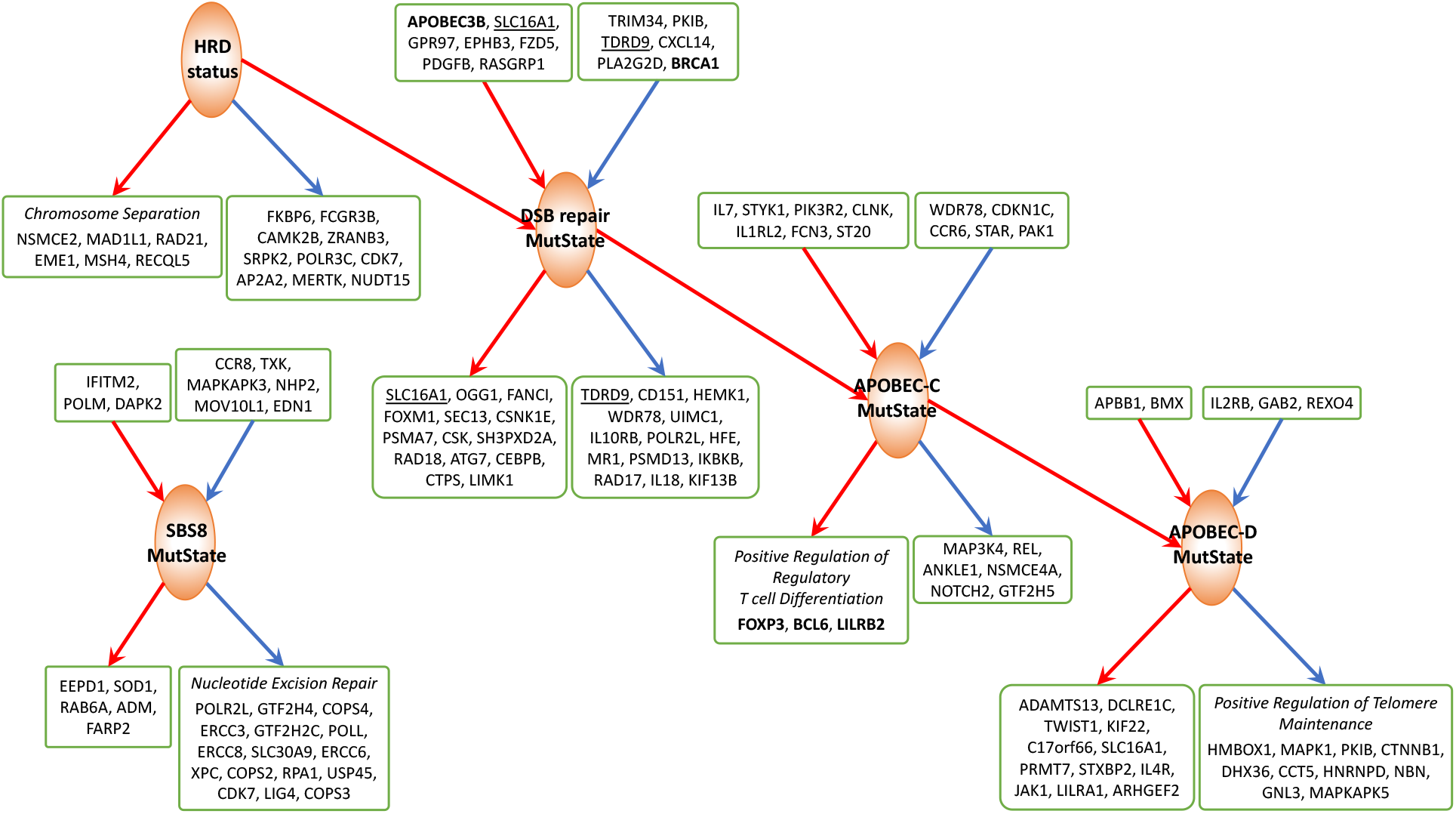
Subnetworks of GSN for BRCA centered on (induced by) MutStates associated with Homologous Recombination, APOBEC and SBS8. Edge and node colors are as in Fig. 1. In boxes, there are the names of the genes adjacent to a given MutState with edge weight cut-off (|*w*_*ij*_| ≥0.01). The genes in bold are discussed in more detail in the text and the genes having bidirected interactions with MutStates are underlined. If the adjacent genes are enriched with specific GO pathways (*q* −*value* < 0.01) then only the pathway genes are provided in the box. The HRD status dominantly contributes to the mutation strength in the DSB repair MutState. Increased DSB exposure leads to an increase of the exposures of Apobec-C and (indirectly) Apobec-D MutStates. SBS8 mutations are linked to the deficiency of nucleotide excision repair. An extended network including all MutStates and an extended list of genes is provided in Fig. S4 in Supplementary Information.

#### Untangling the interactions between APOBEC and DSB processes

Previous studies speculated that APOBEC related mutational signatures can arise in multiple different scenarios. First, double-strand breaks (DSB) created by the homologous recombination deficiency (HRD) provide mutational opportunities for APOBEC enzymes to act on the ssDNA regions, resulting in clustered APOBEC mutations [36,37,29]. In another scenario, a recent study attributed APOBECmediated hypermutations to the normal activity of mismatch repair which also involves creating ssDNA regions, generating *“*fog*”* APOBEC mutations [28]. The complex interplay between APOBEC activities and other DNA repair mechanisms is yet to be elucidated.

Focusing on the interactions of APOBEC signatures with the other MutStates and genes, we observe that GeneSigNet supports a positive influence of the DSB on APOBEC-C MutState, consistent with the assumption that double-strand breaks provide an opportunity for APOBEC mutations. Additionally, our analysis reveals that the expression level of the APOBEC3B enzyme is associated with the strength of the DSB signature. Indeed, a previous study proposed that APOBEC3 proteins are recruited to DSB sites to participate in the DSB repair process [10]. Thus, DSB contributes to an increase in APOBEC-C strength by two different mechanisms: (i) increased mutation opportunity due to ssDNA created by DSB and (ii) increased mutation probability due to increased APOBEC3B expression. Note that increased APOBEC expression would also increase APOBEC mutations in the *“*fog*”* regions proposed in [28].

On the other hand, the activity of APOBEC-D is positively influenced by APOBEC-C activity, without direct relation to DSB. In fact, GeneSigNet inferred a negative influence from HR status to APOBEC-D MutState, confirming that different mutagenic processes are involved in clustered and dispersed APOBEC mutations (Fig. 6).

#### APOBEC hypermutation activates regulatory T cells – implications for immunotherapy

Interestingly, GO enrichment analysis of the genes associated with APOBEC mutational signatures (genes influenced by APOBEC-C MutState) revealed significant enrichment in positive regulation of regulatory T cells (Tregs) differentiation (Fig. 6). Tregs, a subtype of T cells that suppress the immune response, are important for maintaining cell homeostasis and self-tolerance but can also interfere with anti-tumor immune response [38]. The top three genes (FOXP3, BCL6, and LILRB2) positively influenced by APOBEC-C signature are all related to such inhibitory mechanism to immune response [39,40,41]. FOXP3 is a transcriptional regulator playing a crucial role in the inhibitory function of Tregs. BCL6 is also essential for the stability of Tregs that promotes tumor growth. LILRB2 is a receptor for class I MHC antigens and is involved in the down-regulation of the immune response and the development of immune tolerance.

Our results help to understand a complicated role APOBEC mutagenesis holds for immunotherapy. On one side, patients with cancers displaying a high mutation burden are likely to produce tumor-associated neoantigens (mutated peptides presented at their surface) allowing them to benefit from immunotherapy [42]. In particular, the APOBEC mutational signature was identified as a potential predictive marker for immunotherapy response in some cancers [43,44]. Yet,cells carrying a high mutation burden often develop mechanisms of immune tolerance involving activation of Tregs to protect themselves from the destruction [45,46]. Consequently, observed increased number of Tregs in response to high APOBEC mutations may lead to resistance to immune checkpoint inhibitors [47,48]. Thus, our finding suggests that a combined strategy targeting Tregs in addition to immune checkpoint inhibitors would be most beneficial for a better outcome in APOBEC hypermutated breast cancer tumors.

### 2.3 Analysis of the relations between mutational signatures and molecular pathways in Lung Adenocarcinoma

We next analyzed lung adenocarcinoma (LUAD) data using 466 cancer samples from the TCGA project. The exposure levels of 6 COSMIC mutational signatures (SBS1, 2, 4, 5, 13, and 40) present in the exome sequencing data were integrated with the RNAseq expression data of 2433 genes belonging to the DNA metabolic and immune system processes in Gene Ontology terms to uncover influence between signatures and genes (for details, see Supplementary Section S2.2).

#### GeneSigNet uncovers immune response due to smoking

Two prominent mutational signatures in LUAD, SBS4 and SBS5, are assumed to result from exogenous causes [11]. SBS4 is associated specifically with exposure to cigarette smoking in lungs. SBS5 is known to accompany the smoking signature but it is also present in many other cancer types. Previous studies suggested that cigarette smoking stimulates an inflammatory response [9]. Consistent with these findings, the genes identified by GeneSigNet as influenced by SBS4 and SBS5 MutStates are indeed enriched with immune response genes (Fig. 7).

**Fig. 7.**
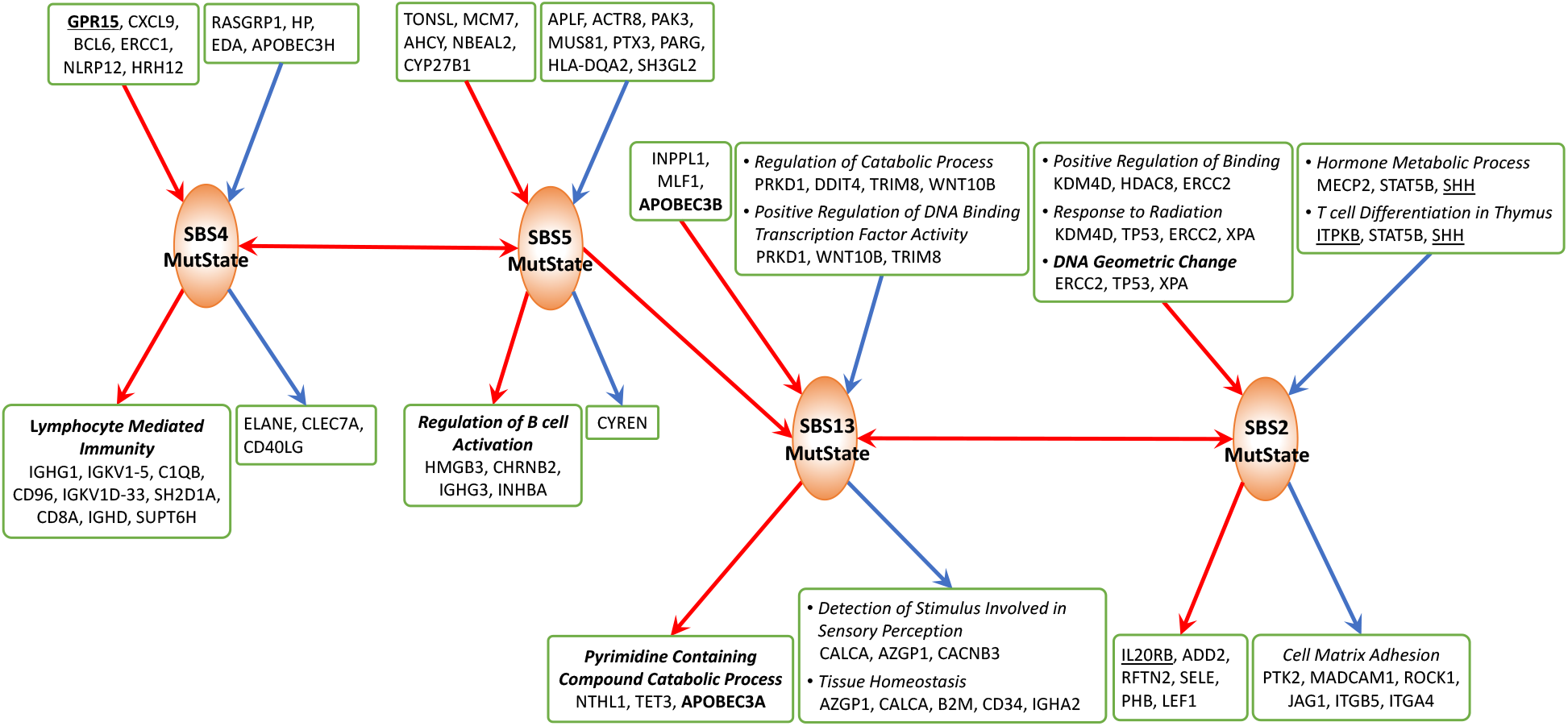
Subnetwork of LUAD GSN centered on MutStates known to be related to smoking and APOBEC. The meaning of colors and boxes is the same as in Fig. 1 and 6. An extended network including all MutStates and an extended list of genes is provided in Fig. S5 in Supplementary Information.

GeneSigNet also identified the influence of the signatures SBS4 and SBS5 on two APOBEC signatures – SBS2 and SBS13. The APOBEC signatures are associated with immune response and this relationship is consistent with the previously proposed immune activation due to smoking exposure [49]. In addition, GeneSigNet correctly captured the association of SBS13 (consequently SBS2) with the expressions of APOBEC3B and APOBEC3A enzymes, and also identified the association of SBS13 with pyrimidine related catabolic processes, potentially reflecting the fact that SBS13 involves a pyrimidine to pyrimidine mutation (Fig. 7).

Finally, tobacco smoking is known to induce GPR15-expressing T cells; although the exact role of GPR15 in response to smoking is yet to be elucidated [50]. Therefore, we investigated whether the results of GeneSigNet provide additional insights into it. Consistently with previous studies, GeneSigNet inferred a strong association between GPR15 and SBS4 (without resolving the direction, see also Fig. S5 in Supplementary Information). Next, we analyzed the influence that GPR15 has on other nodes of the GSN network. The results of GeneSigNet suggest that GPR15 is involved in the negative regulation of several genes related to chemotaxis, including IL10, a cytokine with potent anti-inflammatory properties, and has a positive impact on lymphocyte migration and leukocyte mediated cytotoxicity (Fig. 8).

**Fig. 8.**
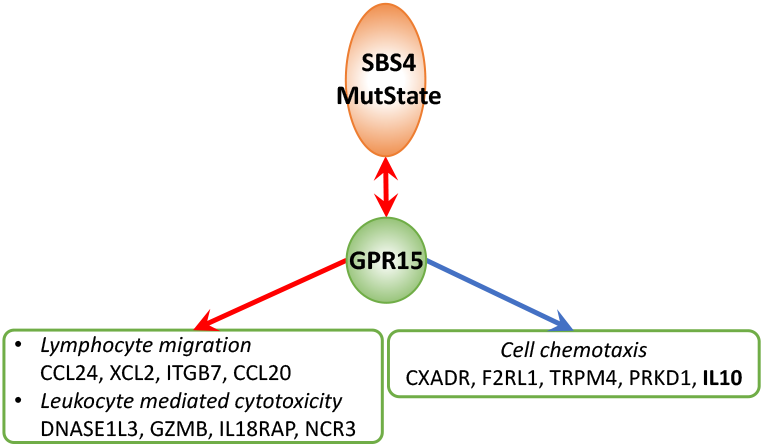
Subnetwork of Lung Adenocarcinoma GSN centered on the node representing GPR15 gene. Expression of the GPR15 gene contributes to the activation of immune responses. The meaning of edge and node colors, and boxes is the same as in Fig. 1 and 6. An extended network including all MutStates and an extended list of genes is provided in Fig. S5 in Supplementary Information

#### GeneSigNet points to the role of DNA geometric changes for APOBEC signature SBS2

As discussed earlier, APOBEC can only act on single-stranded DNA (ssDNA). Interestingly, one of the GO terms associated with SBS2 MutState identified by GeneSigNet is DNA geometric change (Fig. 7). DNA geometric changes are local changes of DNA conformation such as bulky DNA adducts (a type of DNA damage due to exposure to cigarette smoke) or DNA secondary structures such as Z-DNA, cruciform, or quadruplex. Indeed, these structures often involve the formation of ssDNA regions which, in turn, provide mutation opportunities for APOBEC enzymes [51,52,53]. The formation of DNA secondary structures is often associated with DNA supercoiling - a form of DNA stress that is resolved by Topoisomerase 1 (TOP1). Interestingly, GeneSigNet identified a negative influence of TOP1 expression on one of the genes (XPA) contributing to this GO term. This suggests a relation between DNA stress mediated by TOP1 and APOBEC activity.

## 3 Discussion

Elucidating the nature of mutagenic processes and their interactions with cellular processes is of fundamental importance for understating cancer etiology and guiding cancer therapy. Here, we propose GeneSigNet, a new network-based approach that infers the relation between gene expression and the strength of mutation patterns (signature exposures) allowing us to uncover the relations between signatures and processes involved in DNA repair and immune response among other cellular processes. Recognizing the limitations of the previous approaches, GeneSigNet relies on a construction of a sparse directed network. For each node (gene or MutState), it selects a sparse set of incoming edges representing dominating incoming effects so that their combination explains the activity of the node. Aiming to capture the most dominant influences, the method utilizes sparse partial correlation coefficients. In general, the inference of direction of influential relations from statistical dependencies without any support from prior knowledge about regulatory mechanisms is highly challenging. However, it has a potential advantage of capturing the real dependencies in the investigated condition instead of constraining the outcomes subject to the general prior information. GeneSigNet provides an important step toward this direction that is independent of the specific application considered in this study.

Overall the relations discovered by GeneSigNet are consistent with the current knowledge, boosting the confidence in the method*’*s applicability. In addition, GeneSigNet provided several new biological insights concerning the relation between mutagenic processes and other cellular processes. For example, the uncovered relation between APOBEC hypermutation and activation of regulatory T-Cell can have an important implication in immunotherapy. We note that focusing on a sparse set of edges reduces the power of GO enrichment analysis and requires more specific biological knowledge for interpreting the results. Yet, this potential disadvantage is compensated by the compelling mechanistic insights provided by the method.

## Acknowledgements

This study was supported by the Intramural Research Programs of the National Library of Medicine (NLM), National Institutes of Health, USA. We thank Jan Hoinka for his helpful comments on the manuscript.

## Supplementary Information

## S1 Methods

GeneSigNet constructs a sparse network representing the directed dependencies among nodes in a Gene-Signature Network (GSN). The nodes corresponding to genes and MutStates (mutational signatures) are considered random variables and the activities of the variables are denoted as expressions of genes and exposures of mutational signatures across patients (samples). Given an input matrix describing the activities of variables, our network construction method consists of two complementary steps: (i) a sparse partial correlation technique (SPCS) to obtain an initial weighted-directed network by estimating statistical dependencies (edges) between variables (nodes) and (ii) a partial higher moment strategy to refine the initial network by orienting bidirectional edge. A high-level description of GeneSigNet is provided in Section 2.1 in the main manuscript, and the detailed description and related literature are presented below.

### S1.1 A sparse partial correlation selection (SPCS)

In the proposed model, the network inference algorithm does not distinguish between the two types of nodes (genes and MutStates) and infers dependencies between variables based on their observed activities. In this setting, correlation networks are widely used to explore and visualize dependencies in high-dimensional data. However, without assuming prior knowledge, an ordinary correlation itself provides no means to distinguish between influence and affected factors in underlying causal processes [17].

Bayesian networks can be used to infer causal relations of nodes representing their local conditional dependencies via a directed acyclic graph (DAG) [16,17,54,55,56]. Alternative to constructing a DAG, directed partial correlation (DPC) [57] and regression tree based GENIE3 [24] methods have been proposed to uncover conditional dependencies in observed data. However, learning the structure of Bayesian networks from large data is computationally challenging [56] and the inferred networks are always acyclic and thus they do not support feedback loops [58]. On the other hand, DPC and GENIE3 return a complete list of interactions with non-zero weights of connectivity strengths, hence generating fully-connected networks in which the choice of an optimal confidence threshold is left open. Other methods such as the sparse partial correlation estimation (SPACE) [59] and its extension (ESPASE) [60], consider a penalized regression approach to construct the gene regulatory network. Specifically, their joint loss function requires predetermined penalty weights to combine the regression losses over all the response variables. Although the two methods utilize sparse variable selections, both estimations provide symmetric weight matrices representing weighted-undirected networks.

In this work, we modeled directional dependencies as sparse partial correlation coefficients which are obtained by minimizing a least square error subject to the unit *l*_1_ norm ball. Inspired by the theoretical foundations for approximating partial correlations (Box 1), SPCS selects the best combination of a small number of explanatory factors that, under the conditional dependency assumption, explains the activity of each node in the GSN (Fig. 2B).

BOX 1: PARTIAL CORRELATIONS CAN BE APPROXIMATED BY MULTIVARIATE REGRESSION COEFFICIENTS

Consider the ordinary correlation *ρ*_12_ between two random variables *ν*_1_ and *ν*_2_. If *ν*_1_ and *ν*_2_ are correlated with *n* − 2 other variables *ν*_3_, *ν*_4_, …, *ν*_*n*_, we may regard *ρ*_12_ as a mixture of a direct correlation between *ν*_1_ and *ν*_2_ and an indirect portion due to the presence of other variables correlating with *ν*_1_ and *ν*_2_. The partial correlation measuring the direct portion of the total correlation can be represented as a correlation between *ν*_1_ and *ν*_2_ after removing effects due to other variables by a linear regression and the least square linear regression coefficients are proportional to the partial correlation coefficients [21].

The entire network is represented as a weighted-directed graph, 𝔾 = (𝕍, 𝔼), where a set of nodes 𝕍 represents genes and MutStates, and a set of edges 𝔼 represents the relationships among the nodes. As mentioned in Section 2.1 the network inference algorithm does not distinguish between the two types of nodes and assumes the nodes as random variables.

Let 𝕀 denote the index set of *n* variables representing the nodes in 𝕍. The nodes have observational activities over samples, and a *p × n* matrix *X* =*{x*_*ij*_ *}* represents the data consisting of expressions of *m* genes and exposures of *n* −*m* mutational signatures across *p* samples (patients). Assuming the incoming effects on a given variable *f* from its dominating covariates, the observed value *x*_*if*_, corresponding to *i*-th sample, can be approximated as the following affine combination

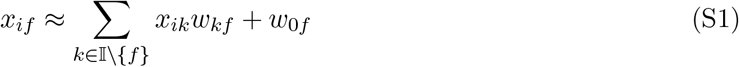

where 𝕀 \ {*f*} denotes the index set of all variables except for the response variable *f*, and *w*_*\*f*_ ∈ *R*^*n*−1^ denotes the weights of incoming effects on *f* from the other *n*− 1 variables and *w*_0*f*_ denotes the intercept adjusting the fitness between the response variable and its prediction.

Thus, our goal is to find the minimum of the least square error function subject to a unit *l*_1_ norm constraint on *w*_*\*f*_ as following

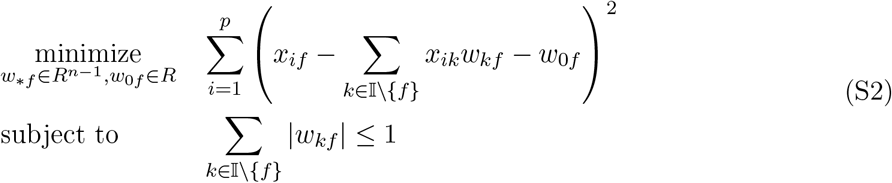

For a response node (affected) *f*, a vector *w*_*\*f*_ = (*w*_1*f*_, *w*_2*f*_, … *w*_*f*−1*f*_, *w*_*f*+1*f*_, … *w*_*nf*_) ^*T*^, the solution to the problem in Equation (S2), represents the weights of incoming effects from the other *n* −1 nodes (influence factors) in the network. The *l*_1_ norm constraint pushes the weights of insignificant effects towards zero and allows selecting only dominant influences on the given response variable. Hence, this regularization acts to avoid over-fitting issues. A non-zero *w*_*kf*_ (*k* ≠ *f*), selected for the node *f*, denotes the partial correlation coefficient representing the potential effect of the node *k* on the node *f*. In this setting, focusing on the activity of every single node in the network, the optimization problem in Equation (S2) considers all possible combinations of incoming effects from the other *n* − 1 nodes and selects the best combination with their optimal influence weights to explain the given response activity under conditional dependency assumption. Therefore, the directed relationship between nodes *k* and *f* is estimated in the presence of the other *n* − 2 variables.

### S1.2 Solving SPCS model

An accurate solution of the problem in Equation (S2) is critical for the robust estimation of dependencies in the GSN. Although the least square error function is convex, the *l*_1_ norm sparsity constraint is non-differentiable and derivative-based techniques such as Lagrange multipliers and Karush-Kuhn-Tucker (KKT) conditions are not directly applicable here due to the non-smoothness. Another attempt to resolve such an issue is to decompose the inequality of the *l*_1_ norm into 2^*n*^ inequality constraints [61]. However, biological networks are often large in scale and it is practically difficult to accurately minimize such a large scale objective function over the exponential number of constraints within a reasonable time. Thus, to approximate the non-smooth constrained optimization, we rewrite the initial formulation in Equation (S2) as an unconstrained form with a penalty term

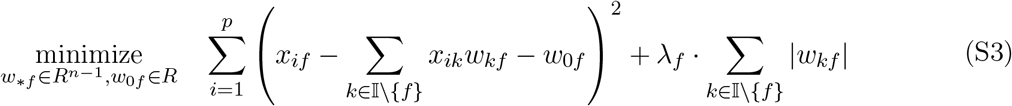

where a tuning parameter λ_*f*_ controls the strength of the penalty term, chosen for the given variable *f* to provide a balance between the least square error term and the *l*_1_ norm constraint in the formulation in Equation (S2). In general, describing the criterion to define a reasonable value of λ_*f*_ is not trivial, due to the incompleteness of information linked to biological relevance. In fact, the exact relationship between the radius of the *l*_1_ norm ball in Equation (S2) and the tuning parameter λ_*f*_ in Equation (S3) is assumed to be data-dependent. Therefore, it is reasonable to use a data-driven strategy for choosing λ_*f*_. Akaike information criterion (AIC) is a statistical technique that provides the relative quality of statistical models for a given data by combining the maximum likelihood estimation of fitness with the number of parameters for inference [62]. The AIC is used in this work to decide the value of λ_*f*_ providing a solution with reasonable total incoming effect on *f* from its dominating factors.

### S1.3 Partial higher moment strategy for influence direction

The solution to the problem in Equation (S2) may provide non-zero weights in both directions for some pairs of variables due to the presence of effects from confounding factors and noise in addition to their real dependencies. This uncertainty may require a complimentary analysis to decide the influence directions of the dependencies.

One way to decide direction of dependency is to perform perturbation experiments [63]. However, optimization of experimental design to predict which combination of perturbations allows to discover influence flows in a given network topology is often challenging and costly [64]. Hence, revealing influence directions by analyzing purely observational data has become a special focus of network biology [65]. Under a confounder-free assumption, higher moment statistics [19] indicate influence direction between two dependent variables from purely observational data. Alternative to the directionality decision for bivariate distributions, a confounder model [66] was recently designed to assign influence directions for several factors under a standard dependency assumption. By combining the key ideas of the two methods, we propose a partial higher moment strategy to propose the influence directions for the bidirected edges in the initial network. The idea is to generate a bivariate distribution for two correlated variables by removing confounding effects from their observed activities, and then decide the influence direction between them using higher moment statistics on the corresponding residual activities (Fig. 2C). Specifically, for two variables having effects on each other (bidirectional edge) in the initial GSN, we first calculate their residual activities by removing effects due to the presence of the other *n* −2 variables [21]. Upon the removal, the corresponding residuals are expected to follow a bivariate distribution and only the dependency between the focused pair remains in their residual activities. Thus, the influence variable can be distinguished from the affected by comparing the higher moments of the two residual distributions. Under the confounder-free assumption, the affected variable is closer to normality than the influence factor and the skewness and kurtosis are the partial higher moment statistics used to measure close-normality of the residual distributions. Under the confounder-free assumption, the affected variable is closer to normality than the influence factor and the skewness and kurtosis are the partial higher moment statistics used to measure close-normality of the residual distributions.

Let *x*_*\*j*_ and *x*_*\*f*_ be *j*-th and *f* -th columns of the given data matrix *X* representing the observed activities of the variables *j* and *f* respectively. Then, the residual activities corresponding to the variables *j* and *f* can be obtained as follows

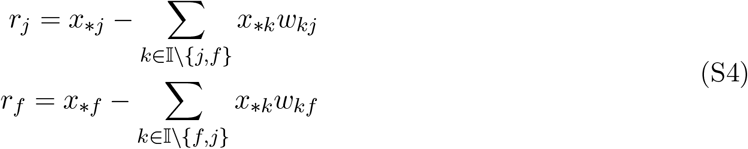

where *r*_*j*_ and *r*_*f*_ are column vectors of size *p*, representing the residual activities after removing the confounding effects from the observed activities *x*_*\*j*_ and *x*_*\*f*_ respectively. For a bivariate distribution of two correlated variables, the affected variable is closer to the normal distribution than the contributing factor, and the higher moment statistics, skewness and kurtosis, can be used to measure the close-normality [19]. As the result of the confounding effect removal, the residual activities *r*_*j*_ and *r*_*f*_ are assumed to follow a bivariate distribution, the influence direction between variables *j* and *f* can be identified by comparing the distribution shapes of *r*_*j*_ and *r*_*f*_. In general, the removal of all confounding effects on a purely observational data is a hard issue due to the possible effects from unobserved latent factors in the domain of genomics. Under this assumption, we use a soft settings for deciding direction between *j* and *f* as following

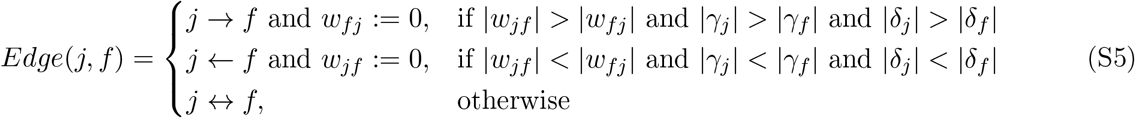

where 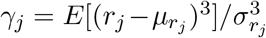 and 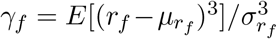 describe the skewnesses of *r*_*j*_ and *r*_*f*_ while 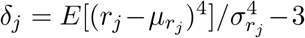 and 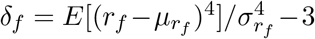 describe the kurtosises respectively, where μ_*·*_ and σ_*·*_ are the mean and standard deviation of the respective variables. That is, as describes in S5, if both moments support the same direction consistent with the stronger weight of the influence then the edge corresponding to the opposite direction is removed. Otherwise, both edges remain in the network. When we first decided dirctionality for every bidirectional edge in the GSN using the strategy S5, the precisions of directionalty inference are 0.597 (*p* −*value* = 7.05*e* −06) in BRCA and 0.559 (*p* −*value* = 4.2*e* −03) in BRCA. Then, we explored different thresholds for edge weight cut-off *τ* to obtain the best set of directed edges. In particular, we accept the direction with a stronger weight if one direction provides a stronger weight than the threshold *τ* while the opposite direction provides a weaker weight compared to *τ*. The edge with the weaker weight is consequently removed from the network. The higher moment-based strategy in Equation (S5) is used to decide the direction if |*w*_*fj*_| ≥*τ* and |*w*_*jf*_| ≥*τ*. The optimal threshold value was chosen (*τ* = 0.0391 for BRCA and *τ* = 0.0521 for LUAD) to maximize the fraction of correct directions (precision) in the set of recovered edges. The performance evaluation is provided in Fig. 5.

Complementary to the partial correlation selection, the partial higher moment strategy provides an influence direction between two correlated variables based on their distribution shapes if the dependency direction is not resolved by the SPCS. Particularly, this strategy is proposed to remove false directed-edges from the initial partial correlation network. To evaluate the robustness of our directionality inference, we tested the reproductivity of the inferred directions by performing a bootstrapping analysis. This analysis was done by repeatedly taking 80% of samples at random without replacement and assigning a direction for each edge using the higher moment-based strategy. Of 3,987 (BRCA) and 2,685 (LUAD) edge directions that were resolved in the presenting results, the reproductivity of each edge direction is tested 100 times during the random sampling. We obtained 83.06% consistent, 2.7% inconsistent and 14.24% unsolved directions for total 3,968 *×* 100 decisions in the bootstrapping analysis on BRCA, and 84.04% consistent, 2.8% inconsistent and 13.17% unsolved directions for 2,685 *×* 100 decisions in the same analysis on LUAD. These results are summarized in Fig. S1.

**Fig. S1.**
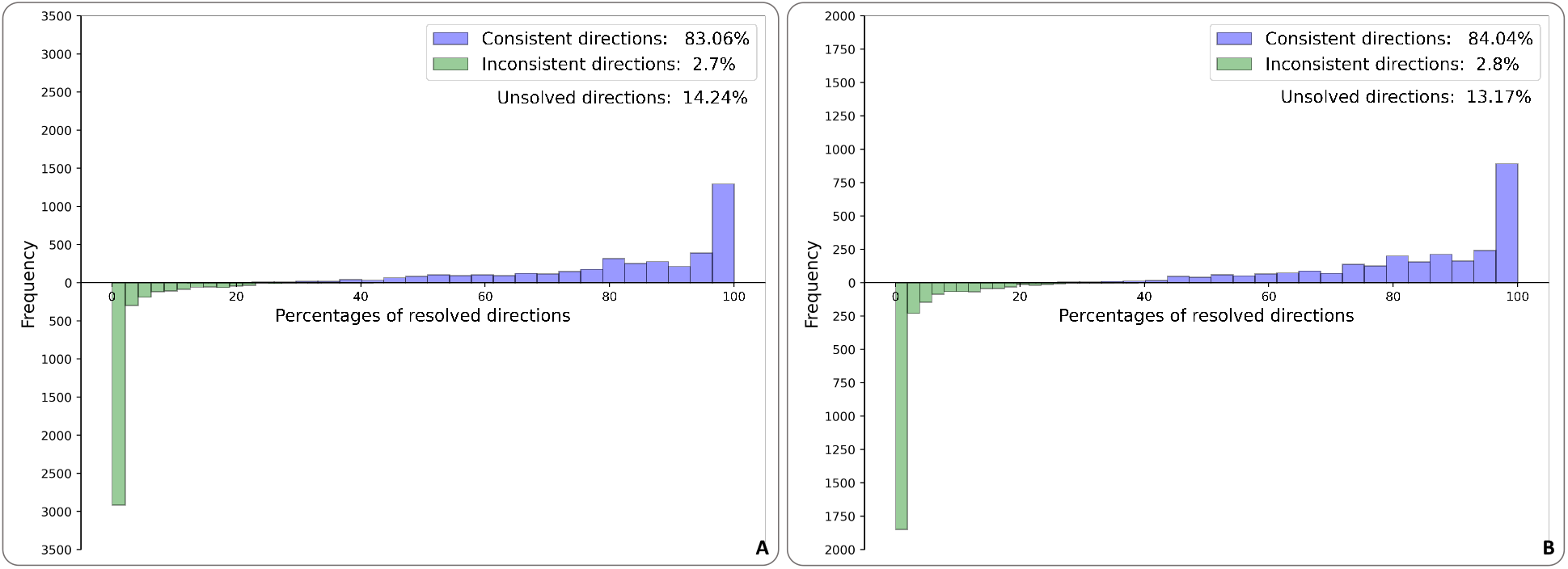
The reproductivity of the inferred directions assigned by the partial higher moment-based strategy. The numbers of consistent and inconsistent assignments during 100 times resampling are shown in the histograms **(A)** BRCA and **(B)** LUAD. The horizontal axis represents the percentages of resolved directions during the resampling while the vertical axis represents the numbers of consistently and inconsistently assigned directions. For example, out of 3,987 edge directions were tested in the bootstrap analysis for BRCA, the consistent percentages of 1,295 directed edges fall in the interval [97%,100%] (the last bin of blue histogram) while the inconsistent percentages of 2,916 directed edges fall in the interval [0%,2%] (the first bin of green histogram). *‘*Consistent directions*’* represents the percentage of the total consistent decisions for overall the directed edges during the 100 times resampling while *‘*Inconsistent directions*’* presents the percentage of the total inconsistent decisions. *‘*Unsolved directions*’* denotes the percentage of the total unsolved decisions that the two higher moment statistics provide contradictory decisions.

### S1.4 Normalization of incoming and outgoing effects

The magnitudes of the effects in the network may provide valuable information to prioritize the candidate associations of genes and MutStates with underlining biological processes because the edge weights denote the contribution scores from influence factors to their affected targets. Hence, it is reasonable to bring the total incoming and outgoing effects of nodes to the same range in the GSN. The total incoming effect on each node was attempted to be normalized into the *l*_1_ norm constraint in Equation (S2). However, the total outgoing effects are free from normalization. Moreover, an additional update described in Equation (S5) was performed to remove edges from the initial network obtained by the SPCS model in Equation (S2).

We adopted a matrix normalization technique, alternate scaling [23], to rescale columns and rows of the weight matrix *W* into the unit *l*_1_ norm ball. This procedure begins with rows in which each is mapped into the unit ball. Then do the same operation on columns, then on rows, and so on, until the sequence of matrices converges. The absolute difference of two consequence updates, by rows (*W*_*rows*_) and by columns (*W*_*columns*_), is used as the convergence criterion such that *‖W*_*rows*_ − *W*_*columns F*_ *‖* < 10^−15^. In the cases of the BRCA and LUAD analysis, the convergence was achieved after only 5 and 6 iterations, respectively. Note that this normalization increases the sparsity of the GSN since every column and row of *W* is iteratively mapped onto the *l*_1_ norm space (*‖w*_*k\**_*‖*_*l*1_ ≤ 1 for *k*-th row and *‖w*_*\*k l*1_*‖* ≤ 1 for *k*-th column of *W*) which rescales the weights to lower values, even assigning zero weights to weak associations during the iterative procedure.

## S2 Materials

### S2.1 Breast cancer data

The normalized gene expression data for 266 breast cancer (BRCA) patients were downloaded from Table S7 in [67]. Gene expression profiles for 2,204 genes involved in either DNA metabolic or immune response processes of the Gene Ontology (GO) database were selected for the analysis.

For mutational signatures, somatic mutation data were downloaded from the ICGC data portal (https://daco.icgc.org, release 22). The 3,479,652 point mutations were assigned to mutational signatures using SigMa [8]. SigMa divided all mutations into two groups, close-by **C**lustered and **D**ispersed mutations, and assigned each of these mutations to one of 12 COSMIC v2 signatures which were previously identified as active in BRCA (Signatures 1, 2, 3, 5, 6, 8, 13, 17, 18, 20, 26 and 30; https://cancer.sanger.ac.uk/cosmic/signatures). From the signatures classified by SigMa as described above, signature phenotype profiles 1D, 2C/D, 3C/D, 5D, 8C/D, and 13C/D that had exposure levels of at least 10% within each group were selected for further analysis (the numbering refers to the COSMIC signature index and C/D denotes signatures attributed to clustered and dispersed mutations). Examining their correlation patterns among patients, some of the signatures were grouped as follows: Signatures 3C/D and 8D were combined into DSB (double-stranded DNA break repair) related signatures, and Signatures 2C and 13C/D into APOBEC related signatures. The remaining signatures are treated separately, resulting in Signature 1, 2D, 5, APOBEC, DSB. A log transformation was consequently performed on exposures of each signature to make its distribution shape closer to a bell curve of normality.

Furthermore, we included binary information of homologous recombination deficiency as an additional variable in the analysis. The binary alteration information was obtained by aggregating functional inactivation information for BRCA1/BRCA2 and 16 other HR genes as provided in Supplementary Tables 4a and 4b of Davies *et al*. [30]. The positive entries were assigned a real value of 4.218 in the SPCS model (Section S2 in Supplementary Information) with the hyperparameter search for the best performance in terms of the means of minimum least square errors and maximum Pearson correlation between responses and predictions over all nodes.

### S2.2 Lung adenocarcinoma data

The expression data (RNA-seq) of the lung adenocarcinoma (LUAD) from The Cancer Genome Atlas (TCGA) project were downloaded from the Genomic Data Commons Data Portal on 2020-06-05 (https://portal.gdc.cancer.gov/). Normalization and variance-stabilizing transformation (vst) of HTSeq count data were performed using DESeq2. Tumor and normal samples were split into different groups and only one sample per donor was kept in each group.

The TCGA LUAD exome mutation spectra were downloaded from Synapse (accession number: syn11801889) and decomposed into COSMIC v3 signatures SBS1, SBS2, SBS4, SBS5, SBS13, SBS40, and SBS45 using the quadratic programming (QP) approach available in the R package SignatureEstimation [68]. Only signatures predominantly active in lung cancer (signatures that were present in at least 5% of samples and were responsible for at least 1% of mutations) were considered based on the initial sample decomposition provided by Alexandrov et al. [2] (Synapse accession number: syn11804065). Signature SBS45 is likely a sequencing artifact so it was omitted from further analyses presented in this study. The same log transformation used in BRCA analysis was performed on signature exposure data as well.

We analyzed 466 tumor samples that had both gene expression and mutational signature exposure data available. We analyzed 2,433 genes belonging to the DNA metabolic process and immune system process in GO terms (genes that are not expressed in at least 10% of the samples were omitted). The gene expression and mutational signature exposure data were combined to form an input data matrix.

### S2.3 Schema for generating observational data

To compare the performance of GeneSigNet with Regression tree [24], GeneNet [17] and LiNGAM [16] methods, we further implemented the data simulation schema provided in LiNGAM [16] to generate a data set for dependent variables. The assumption is that the observed data was generated from a process with the properties that the variables *ν*_*k*_, *k* ∈ *{*1, …, *p}* are arranged in an influence order, such that no later variable influences in any early variables and the value assigned to each variable *ν*_*k*_ is a linear function of the values assigned already to the earlier variables, plus a noise term *e*_*k*_ (external influence) drawn from non-Gaussian distributions as following equation

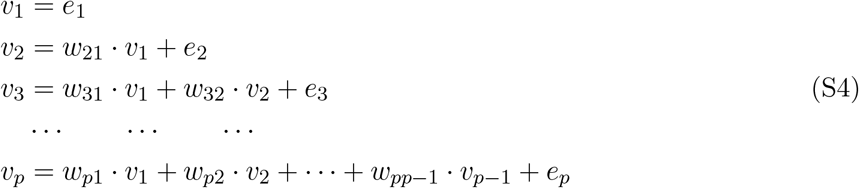

where *w*_*ij*_ denotes the weight of effect from a variable *ν*_*j*_ on its target variable *ν*_*i*_. The recursive process (S4) can be represented graphically by a directed acyclic graph (DAG) where nodes are ordered in such way that no node has an edge that points to a node earlier in the order. This also implies that the adjacency matrix has only zeros on and above the diagonal. For generating a sparse DAG, non-zero weights *w*_*ij*_ were randomly scattered on the strictly lower triangular part of a zero matrix and the row index of the matrix denotes the influence other of the variables in the data generation process. That is, the samples of the seed variables in the order were first generated from standard normal distribution, then subsequently passing their influences through their down stream variables in the directed path to generate samples for the other variables.

## S3 Supplementary Figures

**Fig. S2.**
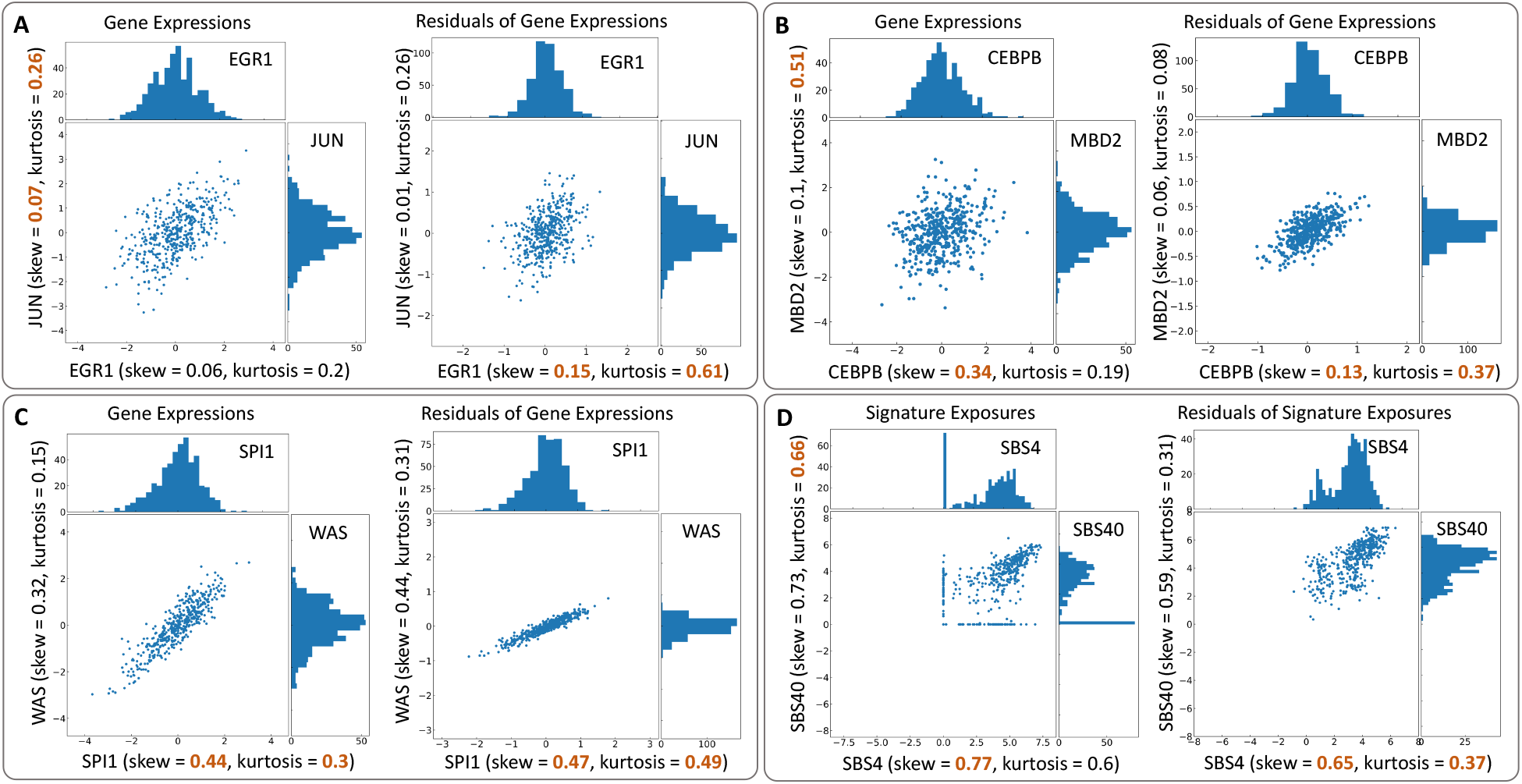
The removal of confounding effects allows the higher moment statistics to distinguish influence factor from its own target. For two random variables in each panel, the joint distribution of their observational activities is shown on the left while that of corresponding residual activities is on the right. The experimentally confirmed regulatory relations [22] indicate the correct directions. The values in brown highlight the influence factors. **(A)** Without removing confounding effects from expressions of EGR1 and JUN, both moments provided an incorrect direction. After removing the confounding effects, the two moments indicate the correct direction. **(B)** Before removing the effects, the skewness provide the direction from CEBPB to MBD2 while the kurtosis supports the opposite direction. However, the proposed higher moment-based strategy indicates the correct direction **(C)** Either without or with the effect removal, the two moments provide the experimentally confirmed influence in WAS from SPI1. Although the higher moment-based strategy did not suggest any changes on the directionality decision, the effect removal uncovers a stronger association between the two genes. **(D)** The two moments provide contradictory decisions without removing the confounding effects from the exposures of the two mutational signatures. However, with the removal, the inferred direction is from SBS4 to SBS40.

**Fig. S3.**
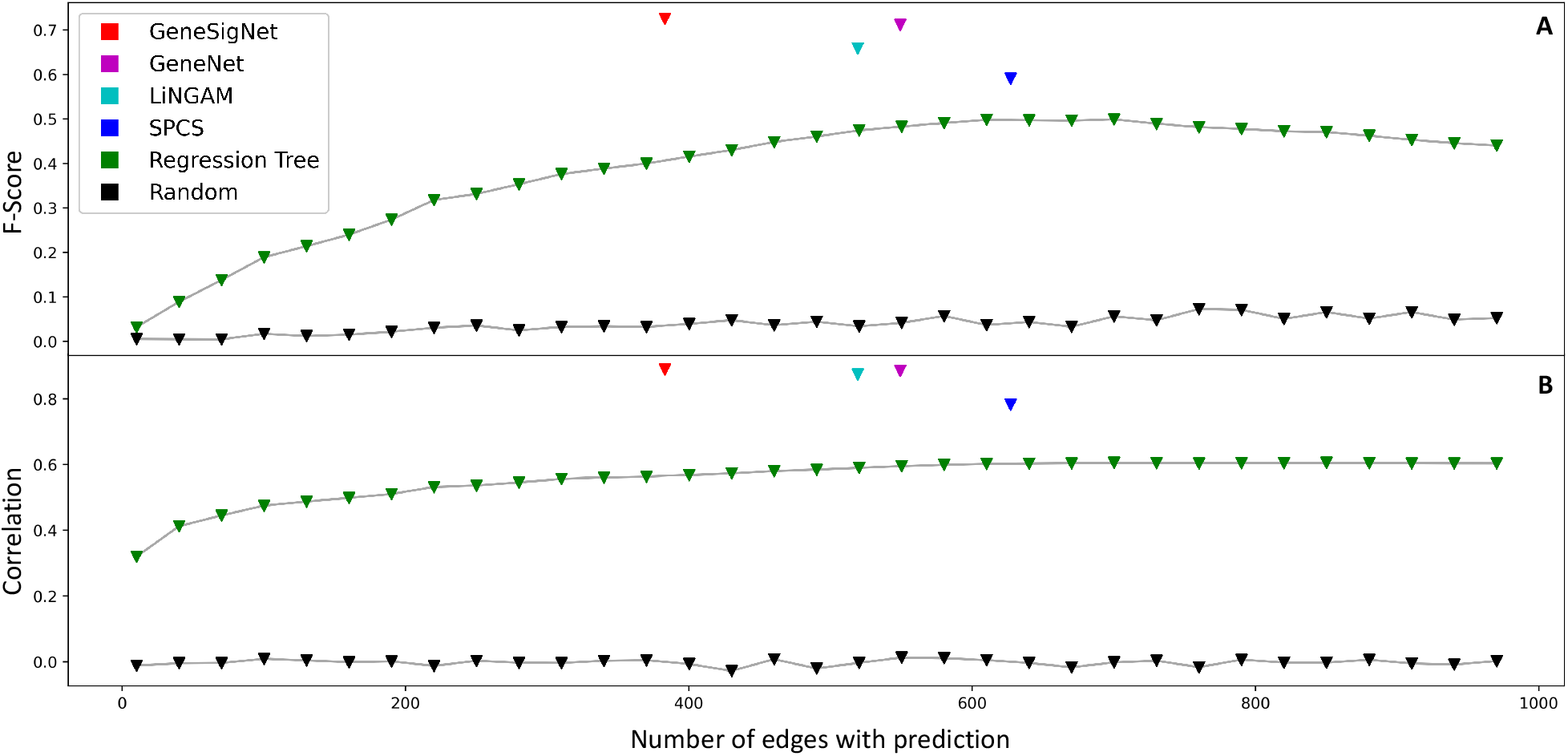
The horizontal axis denotes the number of edges with prediction. The F-score represents the prediction accuracy (A) and the correlation between generated and estimated weight matrices represents the reproductivity of the prediction (B). For Regression tree, absolute values of entries in the reference matrix were used to compute the correlation since this method was designed to provide only positive influence weights between random variables.

**Fig. S4.**
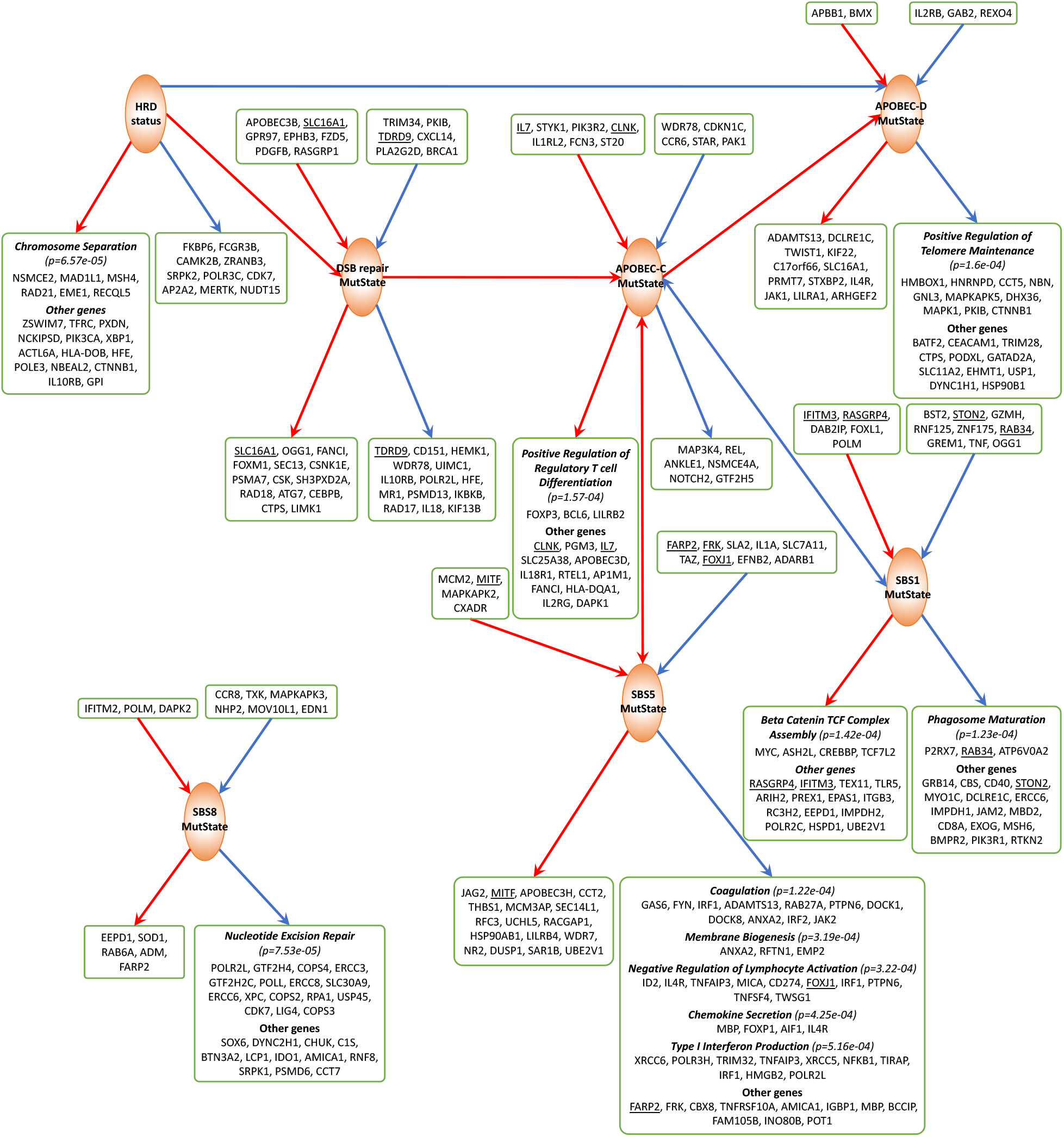
Subnetworks of GSN for BRCA centered on MutStates. Edge and node colors are as in Fig. 1. If the genes adjacent to a given MutState are enriched with specific GO pathways (*q* −*value* < 0.01), the names of the enriched genes are provided under the pathway names in a box and the other adjacent genes with edge weight cut-off (| *w*_*ij*_| ≥ 0.01) are titled as *‘*Other genes*’* in the same box. If the adjacent genes are not enriched with any specific GO pathways (*q* −*value* < 0.01), the names of the genes with edge weight cut-off (|*w*_*ij*_| ≥0.01) are provided in a box. The complete lists of upstream and downstream sets for MutStates and their enriched GO terms obtained in the analysis on the BRCA data are available as a resource at the GeneSigNet GitHub repository (https://github.com/ncbi/GeneSigNet).

**Fig. S5.**
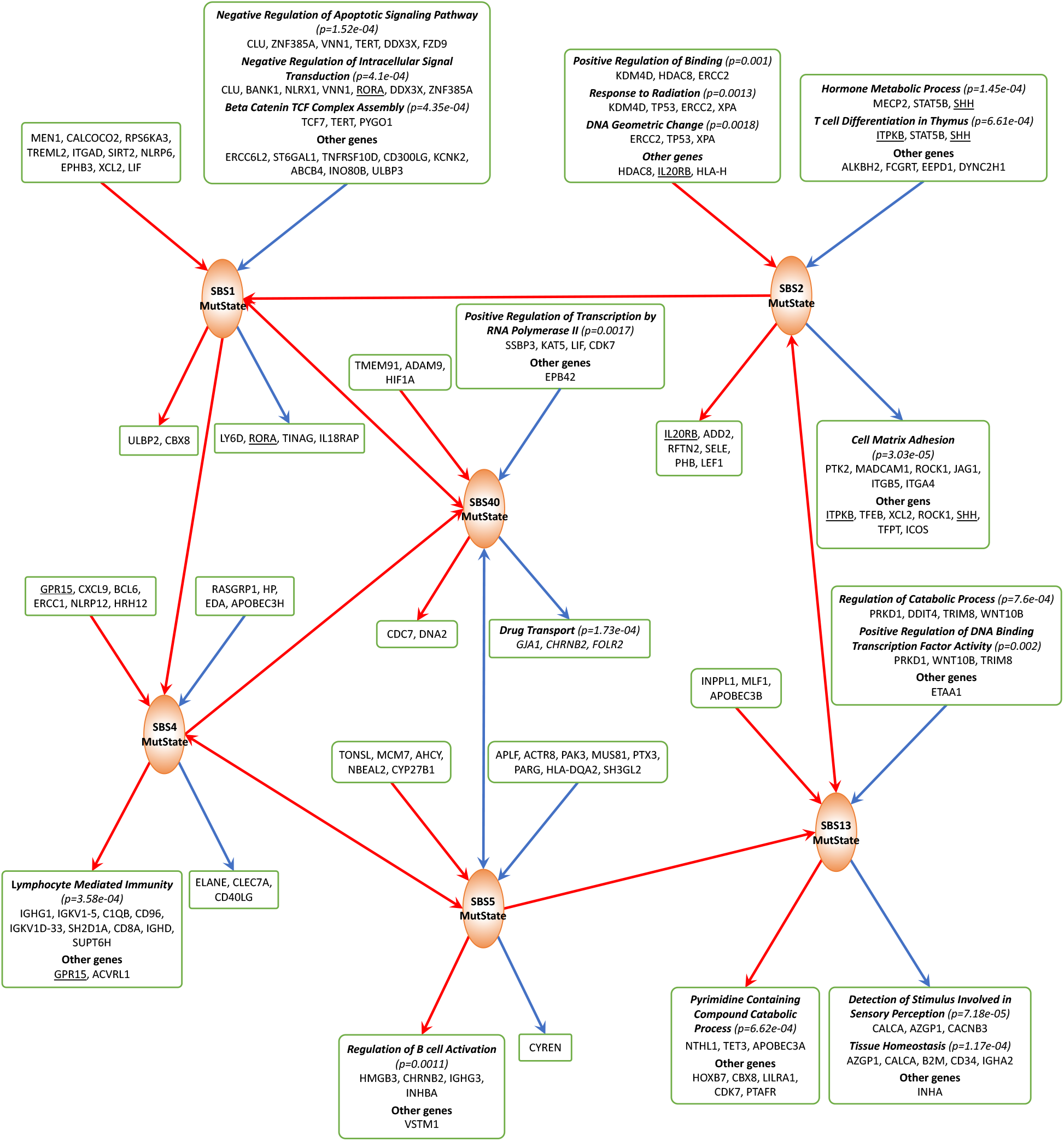
Subnetworks of GSN for LUAD centered on MutStates. The meaning of edge and node colors, and boxes is the same as in Fig. S4. The complete lists of upstream and downstream sets for MutStates and their enriched GO terms obtained in the analysis on the LUAD data are available as a resource at the GeneSigNet GitHub repository (https://github.com/ncbi/GeneSigNet).

